# Single-molecule Dynamic In-Solution Inhibition Assay: A Method for Full Kinetic Profiling of Drug Candidate Binding to GPCRs in Native Membranes

**DOI:** 10.1101/2021.09.16.460640

**Authors:** Tim Kaminski, Vladimir P. Zhdanov, Fredrik Höök

**Affiliations:** In Singulo AB, Pepparedsleden 1, SE-43183 Mölndal, Sweden; Boreskov Institute of Catalysis, Russian Academy of Sciences, Novosibirsk 630090, Russia; Department of Physics, Chalmers University of Technology, SE-412 96 Gothenburg, Sweden

## Abstract

Kinetic profiling of drug–target interactions using surface-based label-free technologies is well established for water-soluble pharmaceutical targets but is difficult to execute for membrane proteins in general and G-protein–coupled receptors (GPCRs) in particular. That is because surface immobilization of GPCRs tends to alter their configuration and function, leading to low target coverage and non-specific binding. We here describe a novel assay for kinetic profiling of drug binding to the GPCR human beta 2 adrenergic receptor (β_2_AR). The assay involves temporally-resolved imaging of the binding of individual β_2_AR-containing cell membrane-derived liposomes to a surface-immobilized ligand in the presence of screened drugs. This approach allowed to determine association and dissociation constants of β_2_AR and suspended alprenolol (antagonist) and fenoterol (agonist). The set-up combines a 384 well-plate sensor chip with automated liquid handling and the assay takes minutes to complete, making it well adapted for drug screening campaigns.

## Introduction

Surface-sensitive optical biosensing has become a golden standard for quantifying the interaction kinetics between drug candidates and pharmaceutical targets. Technologies based on surface plasmon resonance (SPR), biolayer interferometry (BLI), and optical waveguide grating (OWG) have proven compatible with rapid screening of both small- and large-molecular weight drugs, and provide crucial information in the early pharmaceutical development phases^1^. Conventionally, surface-based bioanalytical sensing technologies are used in direct binding assays, where the pharmaceutical target of interest is immobilized on a sensor surface. Subsequent exposure of the surface immobilized target to a solution containing a ligand or drug candidate of interest, followed by exposing the sensor surface to a pure solution, make it possible to extract the association and dissociation rate constants, *k*_on_ and *k*_off_, as well as the equilibrium dissociation constant, *K*_d_ = *k*_off_/*k*_on_, reflecting the affinity of the interaction. This provides unique advantages compared with solution-based assays [e.g. nuclear magnetic resonance (NMR), isothermal titration calorimetry (ICT), and thermal shift assays], but these technologies also have intrinsic limitations. In particular, despite major advancements^1–5^, direct binding assays cannot be used for reliable kinetic profiling of the interaction of small-molecule drugs with membrane proteins. The main reason behind this limitation is that the function of most cell membrane proteins depends on the native cell membrane environment. While immobilization of purified cell membranes has been proposed^6^, this strategy results in surface coverage of the target that is too low to generate detectable signals, especially for low-molecular weight compounds^7^. Hence, new approaches allowing kinetic profiling of new drugs against and functional investigation of this major class of pharmaceutical targets are needed^8^.

Several means to overcome this limitation have been explored, such as immobilization of detergent-solubilized membrane proteins^6^, reconstitution of detergent-solubilized membrane protein target into a natural lipid environment, such as liposomes^12^, or the so-called lipid nanodisks^9^. However, even in scenarios when reconstitution preserves both structure and function of the membrane protein target, maintaining membrane protein in a functional state when immobilized at the sensor surface for the time periods required by screening applications is not trivial^10^. This is an especially grave concern during drug development, where kinetic profiling is crucial for a successful selection of a small candidate set from a large number of screened compounds^11^.

Among membrane protein targets, G-protein coupled receptors (GPCRs) stand out as notoriously difficult to study^12,13^. Their key role in signalling pathways, expression in the plasma membrane and druggability are crucial factors that make them the most important protein family to target in drug development campaigns^12^. GPCR structure and function are critically dependent on successful preservation of their native environment^14,15^. Major efforts have been invested into stabilization of GPCRs via point mutations to lock them in a stable conformation prior to purification, reconstitution^16^ and surface immobilization^17^. However, since native GPCRs stochastically oscillate between different conformations, which has an impact on their interaction with natural ligands and drugs^18^, locking GPCRs in a single conformation can significantly alter their binding and signalling profile^19,20^. The lack of reliable broadly applicable biophysical tools, and in particular direct binding assays, to screen the interaction kinetics of small and large molecules with GPCRs in their native environment is a major obstacle for developing drugs targeting GPCRs^21,22^. The importance of filling this need is expected to increase in the future since advances in target identification through modern genomics and gene editing technologies will increase the frequency that previously undrugged or even orphan GPCRs are targeted^23–25^.

Another challenge of using direct binding assays, especially for membrane proteins present in complex environments, such as native membranes, lipids, and/or detergents, is that binding to the target and off-target binding cannot be distinguished. Many drug candidates are hydrophobic, and their non-specific interactions with stabilizing detergents or lipid membranes complicate kinetic profiling. This is commonly addressed by using a reference surface that only displays off-target binding; however, with low signal levels, the design of representative reference surfaces is challenging. Consequently, competition or inhibition-in-solution assays (ISA) are often used as an alternative to direct binding assays. In ISA, a labelled ligand, henceforth referred to as target definition compound (TDC), is designed to bind to a specific site on the target. In the presence of a screened compound, kinetic profiling is obtained by probing the changed rate of TDC binding, using e.g. radioactive labelling or fluorescence resonance energy transfer^26^. Nonetheless, as repeatedly shown,^27–29^ the kinetic window in terms of *k*_on_ and *k*_off_ values that can be quantified is strictly restricted by the kinetic profile of the TDC in relation to that of the screening compound. Further, if the TDC is instead immobilized on a surface with the competition measured using SPR, BLI, or OWG, only *K*_d_ can be determined and TDC has to have a very slow dissociation rate in such studies^30,31^.

To address the challenges associated with kinetic profiling of membrane proteins in general and GPCRs in particular, we have developed and describe here a dynamic ISA (dISA), a novel assay that probes the interaction between membrane protein contained in a fluorescently-labelled lipid vesicle and surface-immobilized TDC. We built on an previous ISA approach^32^ that can only be used for the determination of the equilibrium dissociation constant, *K*_d_. In the present work, full kinetic profiling, that is extraction of *k*_on_, *k*_off_ and *K*_d_,was accomplished by combining total internal reflection fluorescence (TIRF) microscopy with a 384-well microtiter sensor plate, enabling automated liquid injection, rapid mixing in individual microwells (ca. 100 μL volume), and deconvolution of the binding reaction to the sensor surface (Fig. 1). We tested this approach by analysing the kinetics of human β_2_-adrenergic receptor (β_2_AR), an extensively investigated GPCR and major pharmaceutical target for treatment of asthma and hypertension^33,34^, with two established low-molecular weight compounds, the antagonist alprenolol (Mw ca. 249 Da) and agonist fenoterol (Mw ca. 303 Da), with reported affinities in the nM regime^35^. The developed method allows to keep GPCRs in their native environment and provides the same high information content data as state-of-the-art label free biosensing methods such as SPR. This results not only in faster development of biophysical screening assays for GPCRs, but also improves the biological relevance of the data.

**Fig. 1.**
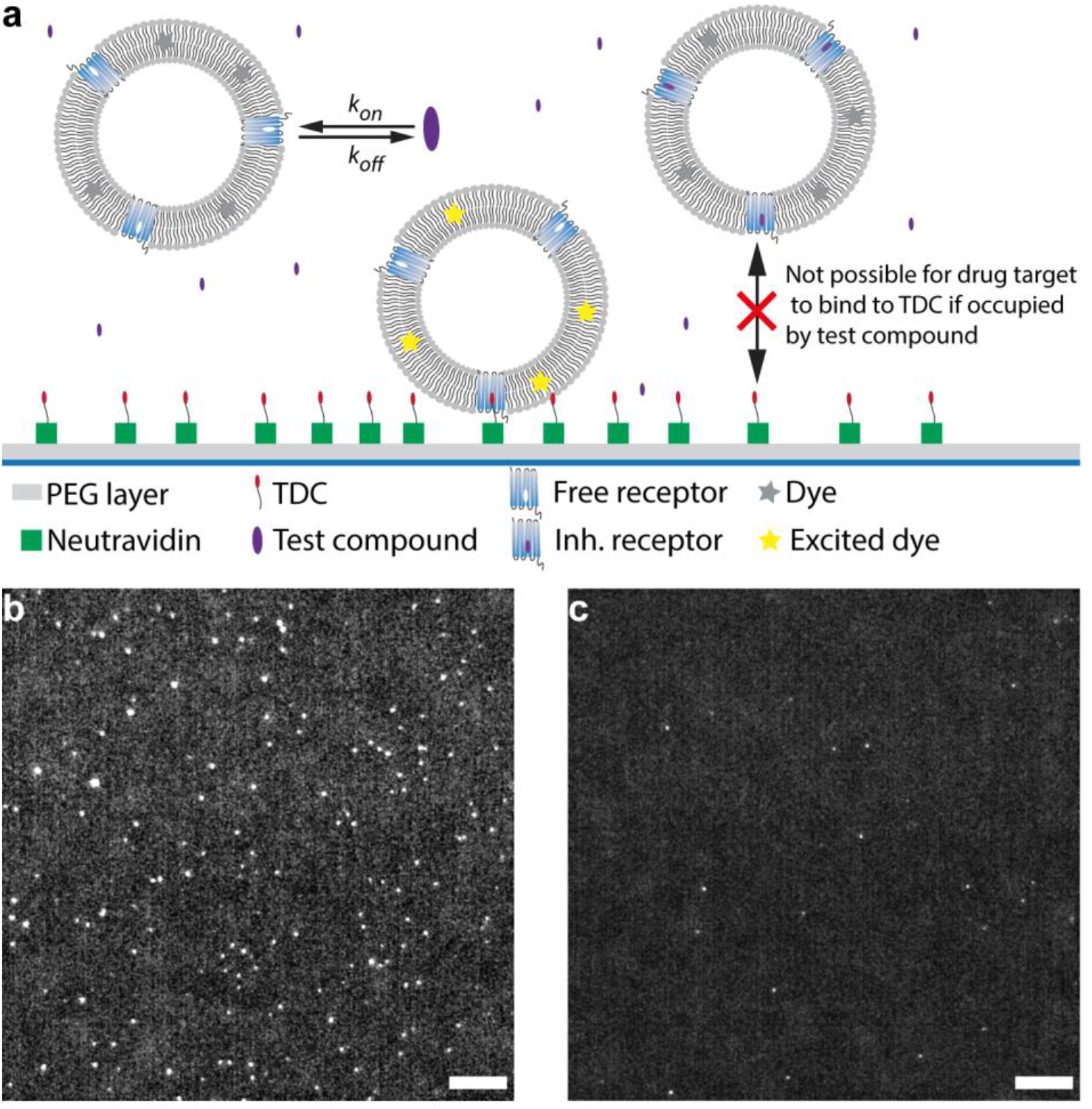
dISA assay developed in the current study. **(a)** In the assay, fluorescently-labelled cell-derived liposomes (CDLs) contain β_2_ adrenergic receptor (β_2_AR). A biotinylated TDC is immobilized via a neutravidin sandwich on the biotin-functionalized PEG-modified surface. Liposomes containing free receptor can bind to the immobilized TDC at the surface. If the receptor is occupied by a test compound, the receptor-containing liposome cannot bind to the surface. Further, the TIRF configuration ensures that only liposomes that are close to the surface are excited and thereby detected. **(b)** Single-frame of a video of β_2_AR CDLs binding to the sensor surface functionalized with alprenolol. Bar, 10 μm. **(c)** As in (b) after preincubation of the same β_2_AR CDLs with 100 nM alprenolol. Bar, 10 μm.

## Results

### Theoretical considerations

The dISA method is inspired by previous work in which surface-sensitive single-molecule microscopy was used to probe how the rate of binding of membrane-protein (target) containing (fluorescently labelled) lipid vesicles to a TDC-modified surface is influenced by the presence of different concentrations of a drug compound directed against the target protein^36^. That way, *K*_d_ of the drug-target interaction can be obtained from the drug compound concentration at which the rate of binding is reduced by 50%, but the approach provides no information about *k*_on_ and *k*_off_ for the interaction between the target and the suspended compound. In the novel dISA method, not only the binding rate of target-containing vesicles to TDC at different compound concentrations is analysed, but also the rate of the transition between two quasi-equilibrium binding states prior to and after the injection of the drug compound of interest. In the following, we describe how this approach enables extraction of both *k*_on_ and *k*_off_ as well as *K*_d_ for the interaction between the target and the suspended compound.

Assuming that each vesicle contains one GPCR that can interact in a reversible manner with the surface-attached TDC, the mean-field approximation in the absence of inhibitor and vesicle diffusion limitations states that the rate of binding can be expressed as

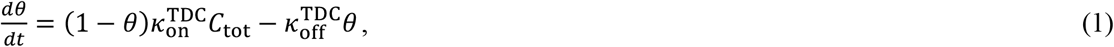

where *C*_tot_ is the vesicle concentration, *θ* is the fraction of TDC compounds occupied by a vesicle, and 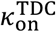 and 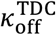 are the attachment and dissociation rates, respectively. Further, 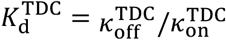 is the corresponding equilibrium dissociation constant. It is also worth noting that the rate of vesicle binding scales linearly with the number of available membrane protein receptors per vesicles in this size regime of ∼100 nm vesicles diameter^37^. Hence, the presence of multiple protein receptors (per vesicle) can be accounted for by including the corresponding scaling into 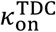. Thus, although the derivation below is strictly valid under the assumption of one membrane protein per vesicle, the presented extraction of the rate constants holds true also if there are multiple membrane proteins per vesicle^37^.

Upon addition of a compound that inhibits the GPCR-mediated vesicle binding to the tool compound, the vesicle concentration, *C*_free_, available for binding to the tool compound is given by

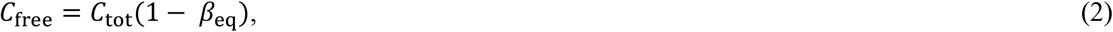

where

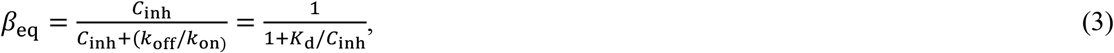

is the fraction of GPCR-containing vesicles occupied by an inhibitor at steady-state, *C*_inh_(≫ *C*_tot_) is the concentration of the inhibiting compound, *k*_on_ and *k*_off_ the association and dissociation rate constants characterizing the interaction between the inhibiting compound and the membrane protein, and *K*_d_ = *k*_off_/*k*_on_ is the corresponding equilibrium dissociation constant.

By operating at low vesicle coverage (*θ* ≪ 1), and by tracking binding events only, Eq. 1 is converted to the steady-state expressions for *t* < *t*_inh_

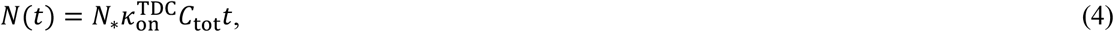

and for *t* > *t*_inh_

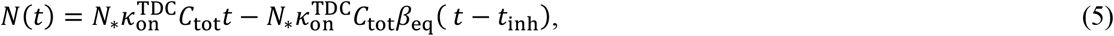

where *N*_*_ is the number of TDCs in the observation area, and *t*_inh_ represents the point in time at which the inhibiting compound is added to the solution. Hence, when *C*_inh_ is known, and by tracking the rate of binding i) before the addition of the inhibiting compound (Eq. 4) and ii) at steady state after complete inhibition (Eq. 5), *K*_d_ can be obtained from the ratio between the slopes measured in the presence of different *C*_inh_.

Further, the rate of the transition between two such equilibria can be obtained from the temporal evolution of the fraction of occupied membrane proteins, *β*, at *t* > *t*_inh_, which is described by

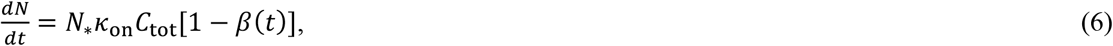

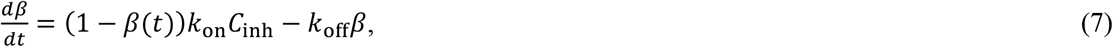

After integration of the latter equation, we have

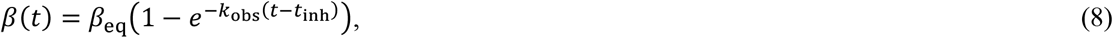

where

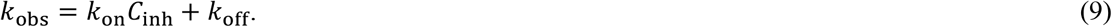

Thus, the temporal evolution *N*(*t*) at *t* > *t*_inh_ during transition between the two equilibria can be expressed by integration of Eqs. 6 and 8, which yields

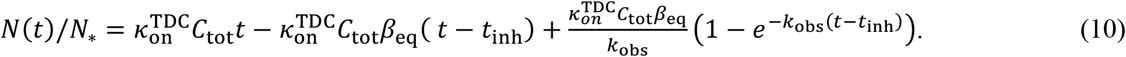

Thus, *k*_obs_ can be directly obtained by fitting the transition curve to Eq. 10. Further, with *K*_d_ known from Eqs. 4 and 5, both *k*_on_ and *k*_off_ can be determined from *k*_obs_.

Concerning the conditions of the applicability of the equations above, we note that their derivation implies that i) the motion of solution after introduction of inhibitor rapidly relax and ii) the diffusion-limited corrections in description of attachment of vesicle to the sensor surface during the time period of the experiment are negligible. The corresponding conditions are given and discussed in Section 1 in the Supporting Information. Practically, one must also include a correction associated with the slight dilution caused by adding to the total sample volume, *V*_tot_, a small volume, *V*_inh_, containing the inhibiting compound: *V*_tot_/(*V*_tot_ + *V*_inh_).

In contrast to the here described single molecule-based method, classical ensemble-based methods such as SPR, BLI, OWG, radioligand binding etc. cannot distinguish the association and dissociation reaction to the TDC. This limitation complicates the analysis of binding kinetics as explained below. In a corresponding ensemble-based ISA experiment, the temporal evolution of the net change in TDC occupancy *θ*(*t*) at *t* > *t*_inh_ is given by^26,38^

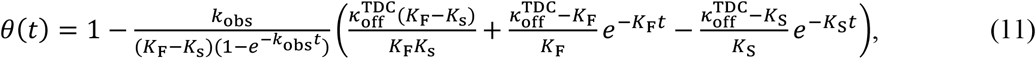

where

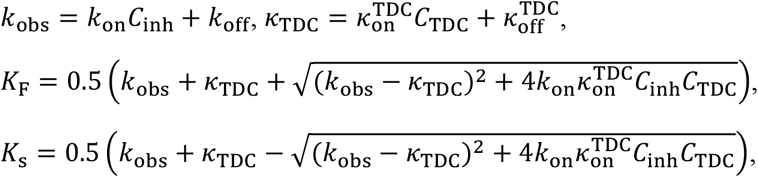

and where *C*_TDC_ is the concentration of suspended TDC. Although Eq. 11 was derived for inhibition in solution, i.e. not for surface-based ensemble averaging methods, a comparison of Eqs. 10 and 11 illustrates that the transition rate in the former depends on *k*_obs_, i.e. *k*_on_ and *k*_off_ only (see Eq. 9), while the latter denotes a complex relationship between and depends on *k*_on_, *k*_off_, 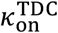, and 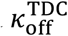. This complication is inherent to all ensemble-averaging ISAs.

### Preparation and characterization of GPCR-containing cell membrane-derived liposomes

To verify the applicability of the devised method for screening of low-molecular weight compounds targeting GPCRs, we selected the human β_2_AR, a well-studied prototypical GPCR^33,34^. Importantly, kinetic data from direct binding assays exist for this protein^39^, allowing direct comparison with the data obtained using the novel assay. To generate β_2_AR-containing liposomes with the highest possible physiological accuracy, we stably transfected CHO-K1 cells with a plasmid coding for a human β_2_AR with a C-terminal fusion with green fluorescent protein (GFP). We identified and selected stable high expression clones based on their GFP-fluorescence. To produce fluorescently labelled cell derived vesicles, as a first step the membrane of adherent cells was stained. The cell membrane was rapidly stained by both DiD and DiI, which are well characterized membrane integrating dyes (Fig. 2a). Upon incubation of fluorescently labelled cells with cytochalasin B, a cell-permeable mycotoxin that inhibits actin polymerization, the cells shed plasma membrane yielding cell-derived vesicles (CDV), with diameters ranging from <1 μm to 10 μm (Fig. 2b), in agreement with previous reports^40,41^. Although neither β_2_AR nor DiD was homogenously distributed in the CDVs (Fig. 2b), co-localization analysis revealed a considerable overlap of DiD and β_2_AR-GFP signals within the CDVs before and after extrusion (Fig. 2c and d; Pearson’s R-value of 0.92 and a Li’s intensity correlation quotient (ICQ) of 0.384). By contrast, a considerably less pronounced overlap was observed for DiI (Supplementary Fig. 1; Pearson’s R-value of 0.43 and a Li’s ICQ value of 0.109). We tentatively attributed this to preferential association of the respective fluorophore with different lipid phases^42^, and our experiment thus suggests that the preferential co-localization of DiD with β_2_AR-GFP is related to a preferential association of β_2_AR-GFP with specific lipid environments^43^. This interpretation was consistent with the absence of a functional response for DiI-labelled cell-derived liposomes (CDL) in the dISA assay (not shown), suggesting that DiI was not efficiently incorporated into vesicles containing β_2_AR; by contrast, DiD-labelled CDLs showed a specific response to the sensor surface.

**Fig. 2.**
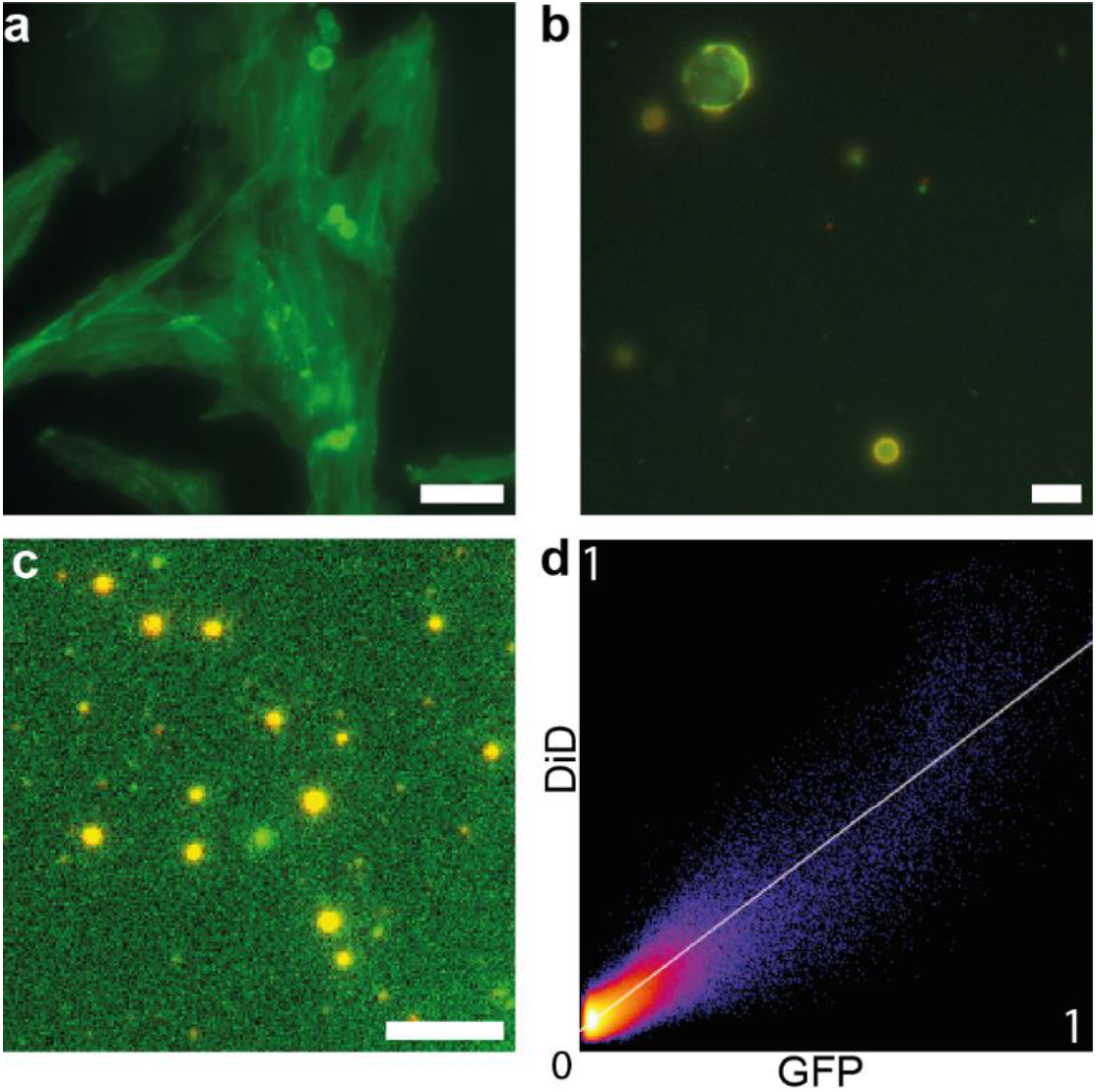
Production of cell-derived liposomes. **(a)** A derived clone of CHO-K1 cells stably expressing the β_2_AR-GFP fusion protein. Bar, 50 μm. (**b)** Co-localized signals of β_2_AR2-GFP (green) and DiD (red) membrane stain in cell-derived vesicles obtained from cells stained with a membrane-inserting dye and incubated with cytochalasin B. Bar, 10 μm. (**c)** Reduced-size vesicles after extrusion through a 100-nm pore membrane. β_2_AR-GFP (green) and DiD (red) fluorescence is shown. Bar, 5 μm. **(d)** 2D histogram of DiD fluorescence plotted against GFP fluorescence of cell-derived liposomes obtained from cells pre-labelled with DiD.

### Full kinetic profiling of the interaction between β_2_AR2-GFP and low-molecular weight compounds

We next fabricated a sensor chip for drug compound screening by passivating a glass surface using a self-assembled biotin-modified co-block polymer^44^, followed by the addition of streptavidin, as described previously^32^, and a subsequent functionalization with the TDC, biotinylated alprenolol. We then explored how the binding rate of β_2_AR2-CDLs to such sensor surface is affected by the addition of known ligands, to finally calculate the full binding kinetics of the respective β_2_AR2 drug compounds.

The key advantage of using single-molecule microscopy to separately identify both the association and dissociation kinetics between the target-containing liposomes and TDC becomes apparent when one compares the total number of vesicles at the sensor surface and the temporal evolution of the number of newly bound vesicles, represented by Eqs. 1 and 10, respectively, as illustrated in Figs. 3a and b for an experiment in which β_2_AR-CDLs are added to TDC-modified sensor surface at *t* = 0, followed by the addition of an inhibiting compound (3 nM alprenolol) at *t* of approximately 20 s. The key difference between these measurements originates from the notion that the number of new binding events (Fig. 3b) identified by single-molecule resolution represents the concentration of suspended liposomes with free ligand binding sites on the β_2_AR2, while the total number of bound liposomes (Fig. 3a) depends both on the duration of the exposure to a certain concentration of free β_2_AR2 -containing liposomes and their dissociation kinetics from the sensor surface, as well as signal disappearance because of photobleaching. Hence, while both approaches can be readily used to extract *K*_d_, if one corrects for photobleaching, kinetic profiling using conventional ISA puts significant restrictions on the properties of TDC (see Eq. 10). By contrast, using dISA, the observed transition rate is not influenced by the binding reaction between the target and TDC, allowing *k*_obs_ to be directly extracted from a fit to the curve in Fig. 3b using Eq. 9.

**Fig. 3.**
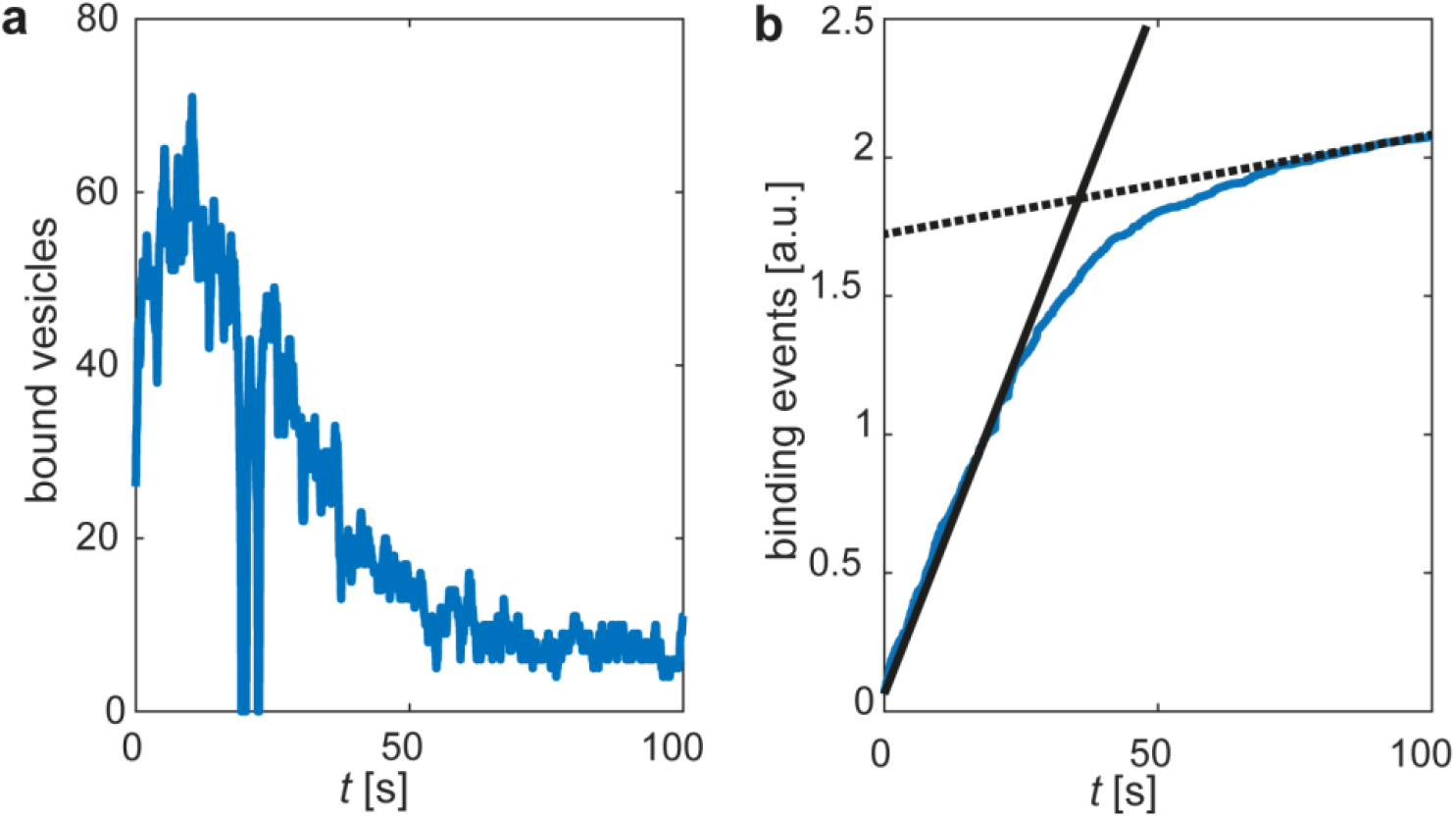
Representative data for signal evolution upon an initial addition of β_2_AR-CDLs to a microwell, followed by the addition of the inhibitor (3 nM alprenolol) at *t* approximately 20 s. **(a)** The total number of bound vesicles at the sensor surface plotted vs. time, in analogy with the signal evolution in ensemble-based sensing that measure the mass concentration at the sensor surface, such as SPR, BLI, or OWG. **(b)** The number of newly bound β_2_AR-CDLs plotted over time, enabling direct resolution of the kinetics of the inhibition reaction. The filled and dashed lines represent the slope before inhibitor addition and the slope after the binding between β_2_AR and the inhibitor reached equilibrium, respectively.

It is also worth emphasising that potential off-target binding sites are often not known *a priori*. To minimize the impact of off-target binding between the surface-immobilized TDC and the target, which typically has short residence time, the measured kinetics can be restricted to events with longer residence time than that of an experimentally determined threshold. This is illustrated by residence time histograms for the interaction between β_2_AR-CDLs and TDC without compound inhibition, and at saturated inhibition for alprenolol (Fig. 4a) and fenoterol (Fig. 4b), a procedure that also efficiently reduces the potential impact from unspecific binding events on the surface. With these specifications, we monitored the rate of the specific binding of suspended β_2_AR-CDL in individual wells, in which the uninhibited response to the sensor surface was first determined. The drug compound was subsequently added to the same microwell, followed by rapid mixing (< 2 s), while continuously monitoring β_2_AR-CDL binding to the surface. The competition between the test compound and TDC for β_2_AR binding resulted in a monotonically decreasing binding rate of β_2_AR-CDLs to the surface-immobilized TDC (Fig. 4c).

**Fig. 4.**
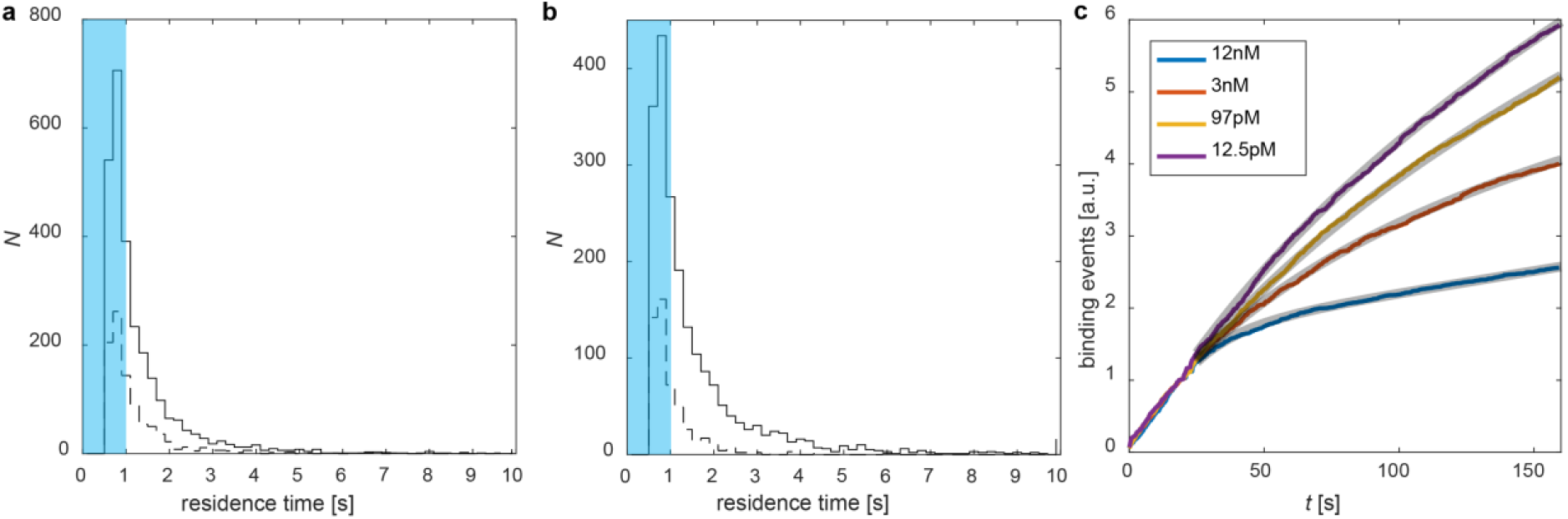
Interaction between β_2_AR-containing cell-derived liposomes (β_2_AR-CDLs) and surface immobilized alprenolol in the absence and presence of small-molecule drug. Residence time distribution of β_2_AR-CDL at a TDC-functionalized sensor surface without prior compound incubation (solid lines) and after saturated compound preincubation (dashed lines) for (**a)** alprenolol (0 and 1 μM) and **(b)** fenoterol (0 and 40 μM). Blue region labels binding events with a residence time <1sec. **(c)** Sensorgrams for β_2_AR-CDL binding to alprenolol–biotin-modified sensor surface and fits of Eq. 10 to the raw data as grey overlays. At *t* = 20 s, different concentrations of alprenolol were added to the microwells.

We subsequently repeated the above type of experiment for a range of compound concentrations above and below *K*_*d*_ of the interaction, allowing determination of *K*_d_, *k*_on_, and *k*_off_ as outlined in the theory section. The analysed two compounds, alprenolol and fenotorol, display orders of magnitude different reported affinities for β_2_AR, in the range of approximately 0.5 nM and 150 nM, respectively^35,45–49^. In good agreement with these reports, we measured *K*_d_ values of 0.11 nM and 119 nM for alprenolol and fenotorol, respectively (Fig. 5a), with the corresponding *k*_on_ values of 1.57 × 10^5^ and 5.4 × 10^4^ M^−1^s^−1^ and *k*_off_ values of 1.6 × 10^−5^ and 6.3 × 10^−3^ s^−1^. The respective *k*_on_ values were extracted from a linear regression of the dependence of *k*_obs_ on *C*_inh_ (Eq. 9 and Figs. 5b, c) while *k*_off_ values were calculated from *k*_off_ = *K*_d_*k*_on_.

**Fig. 5.**
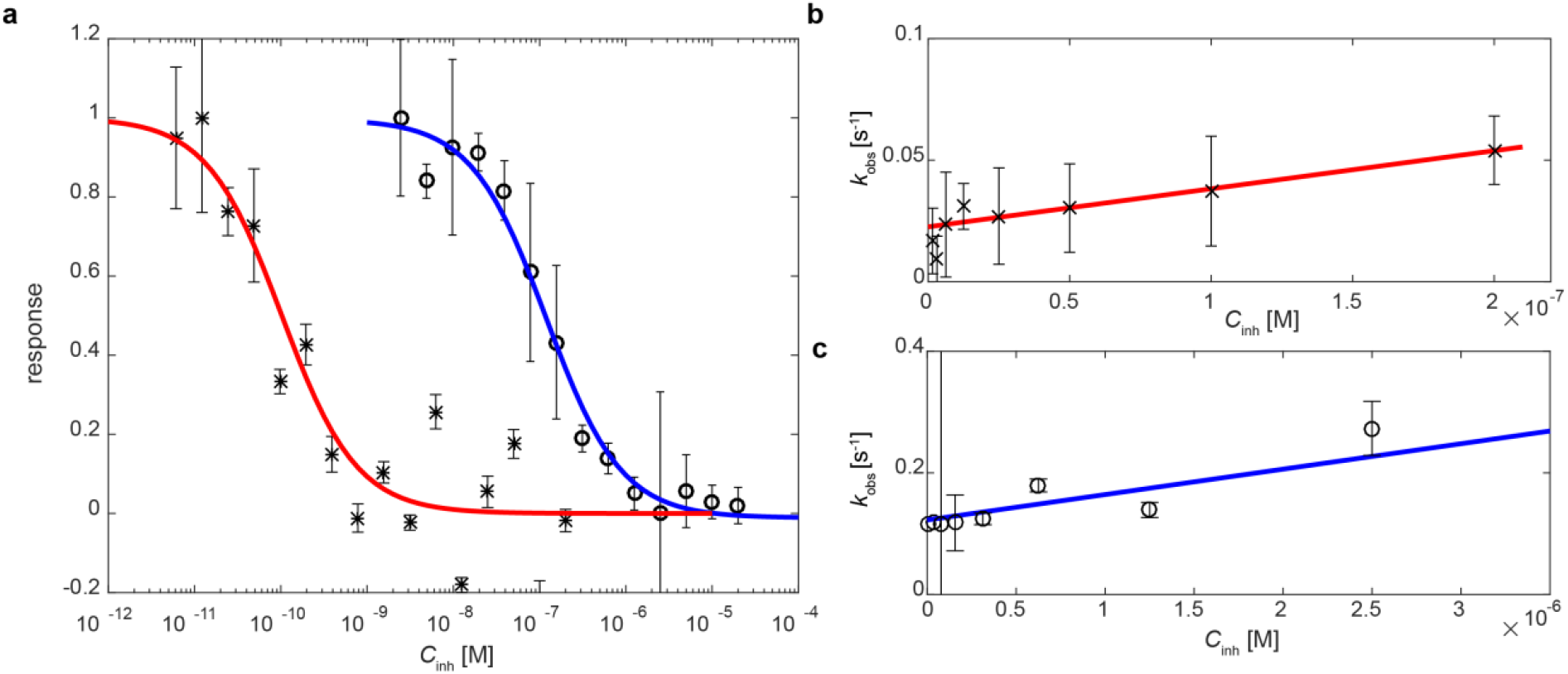
Full kinetic profiling of β_2_AR in cell-derived liposomes. **(a)** Dose-response curve of equilibrium fits using Eqs. 4 and 5 to sensorgrams like those shown in Fig. 4c for the antagonist, alprenolol (×, red), and agonist, fenoterol (o, blue). The response was calculated from the rate of vesicle binding normalized to the uninhibited response and the dose corresponds to *C*_inh_. (**b**) and (**c**) *k*_on_ vs. *C*_inh_ plots obtained from a fit of the sensorgrams like those shown in Fig. 3 using Eq. 9. For all datasets the error bars were calculated via case resampling bootstrapping as explained in the statistical analysis section.

The obtained kinetic profiles were in overall agreement with previous reports using these compounds^35^. Of note, the off-rate constants were up to a factor of 10 slower than those measured using SPR and detergent-solubilized β_2_AR^35^. It is in this context worthwhile to note that for protein stability reasons, the SPR studies were conducted at a relatively low temperature (10 °C), while the current dISA set-up is operated at room temperature (approximately 23 °C) because of the stabilization of β_2_AR by the native cell membrane lipids^14,50,51^. Further, GPCRs are flexible proteins, whose structure fluctuates between different functional conformations. Although previous studies suggest that there is no difference between wild-type β_2_AR and β_2_AR fused to different fluorescent proteins, ^52,53^ we cannot exclude that the fusion of the β_2_AR to GFP impacted the ligand binding kinetics. However, detergents of the type used in previous studies are known to impact the exchange rate between different β_2_AR states^54^ and the different functional states can show different ligand-binding properties^55^. Further, not much is known about the impact of replacing the native membrane environment with detergents on GPCR functionality^14,15^. Hence, even if detergents that allow functional protein analysis are identified^56^, detergents will never perfectly mimic the complex interaction of integral membrane proteins with their native lipid environment^14,18^. For these reasons, we attribute the slower binding kinetics observed here for both compounds to a more native-like conformational flexibility of β_2_AR when analysed in its native membrane than that upon stabilization with detergents. This interpretation is supported by the notion that we conducted the experiments at a higher temperature (23° C versus 10° C) than that reported for detergent-solubilized β_2_AR. In this context, it is also worth mentioning that receptor dimerization is an important aspect of GPCR signalling, which influences ligand and drug compound binding properties of GPCRs^57^. Indeed, it is widely acknowledged that minimalistic experimental models with lower predictive validity lower the success rate of drug development campaigns^58^. The native cell membrane-derived liposomes used in the current study thus provide an attractive platform that likely does not impair receptor dimerization. This could, in turn, open up a more diverse pharmacological landscape for drugs targeting GPCRs than is presently possible^59^. How dISA and its advantages may allow to harness receptor dimerization to develop new drugs will be subject of future studies. Further, slower interaction kinetics improve the ability to detect hits and improve hit differentiation. Hence, the possibility of analysing GPCRs in their native environment with slower, native interaction kinetics is useful for assay configuration, such as off-rate screening, that is often used in fragment-based hit finding strategies^60^.

### General considerations for the selection of TDC compound

In classical Motulsky and Mahan-based competition assays, the measured kinetics of the screened compounds can be influenced by the actual kinetics of the TDC (as described in the theory section) and must therefore be carefully matched with the kinetic profile of the compounds investigated^27,61^. The possibility to restrict the analysis to a defined residence time in a dISA analysis (Fig. 4) puts significantly lower constraints on the TDC, while simultaneously providing a unique opportunity to improve the specificity of the detected signal. In particular, conventional surface-based label-free methods detect signals irrespective of where on the target the binding occurs. By contrast, the method presented in the current study is used to solely detect a signal if the compound binding impacts the actual association between the GPCR and the surface-immobilized TDC. This is particularly beneficial in the context of low-affinity compound screening, such as fragments, since they typically have several binding sites on a protein^62^.

Considering the above, we asked and analysed whether selecting binding events by a defined residence time for the analysis may limit the impact from off-target interactions with the TDC on the extracted kinetics. As in conventional ISA, but in contrast to direct binding assays, dISA benefits from the selectivity provided by the TDC^61^. However, even if the surface is appropriately modified to suppress non-specific binding, any off-target binding between the surface-immobilized TDC and the target-containing vesicles can add complexity to the data, which in turn complicates data analysis^63^. Of note, the off-target binding affinity is usually significantly weaker than the on-target binding, which is typically reflected in higher *κ*_off_ values (shorter residence times). To test how selection of a predefined residence time (1/*κ*_off_) using dISA aids the measurement accuracy, we simulated the dependence of the extraction of the measured equilibrium dissociation constant 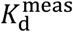 values associated with the screened compound binding to the target on i) the ratio of the equilibrium dissociation constant for the TDC interaction associated with the on-target and off-target binding sites, 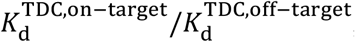, and ii) the ratio of the concentration of on-target and off-target binding sites *C*^on-target^/*C*^off-target^. To do this, we compared the results of simulations without (Fig. 6a) and with (Fig. 6b) residence time-based selection of binding events. Since there are infinite combinations of association and dissociation rates for any 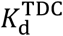, the plots presented in Fig. 6a and 6b were generated assuming that i) changes in *κ*_on_ and *κ*_off_ contribute equally to the changes of dissociation constants 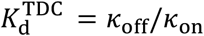, and ii) the test compound has the same affinity, *K*_*d*_, for the on- and off-target sites (see SI for detailed descriptions of the simulations). Figure 6c shows the same mdata as in Fig. 6a and 6b, transformed by plotting 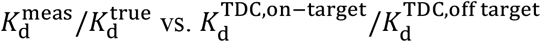 fora selection of *C*^on-target^/*C*^off-target^ ratios. When the concentration of the on- and off-target binding sites is the same, the ratio between 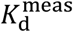 and the true *K*_*d*_ is maximal when the equilibrium dissociation constant for the target interaction with TDC, 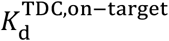, is approximately a factor of _d_ 10 lower than the corresponding value for the off-target interaction, 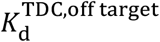, reaching a maximum 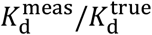 of 2.1 (Fig. 6c). As shown in Fig. 6b, the maximum impact on 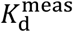, which also in this case occurs at 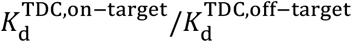 of approximately 10, is reduced to 1.3 by introducing a minimum residence time for binding events selected for analysis (see Fig. 6 legend and SI for the definition of the selection criteria). Further, the advantage of using residence time selection for the extraction of 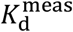 is significant over a wide concentration and specificity range for the on- and off-target binding of TDC, resulting in less stringent specificity requirements for TDC. Indeed, even if the concentration of the off-target binding sites for TDC exceeds that of the on-target binding sites (*C*^on-target^/*C*^off-target^<1), accurate *K*_d_ determination is still feasible (Fig. 6b and 6c). However, kinetic selection of binding events also reduces the number of the detected binding events, which means that careful selection and optimization is required to achieve optimal results.

**Fig. 6.**
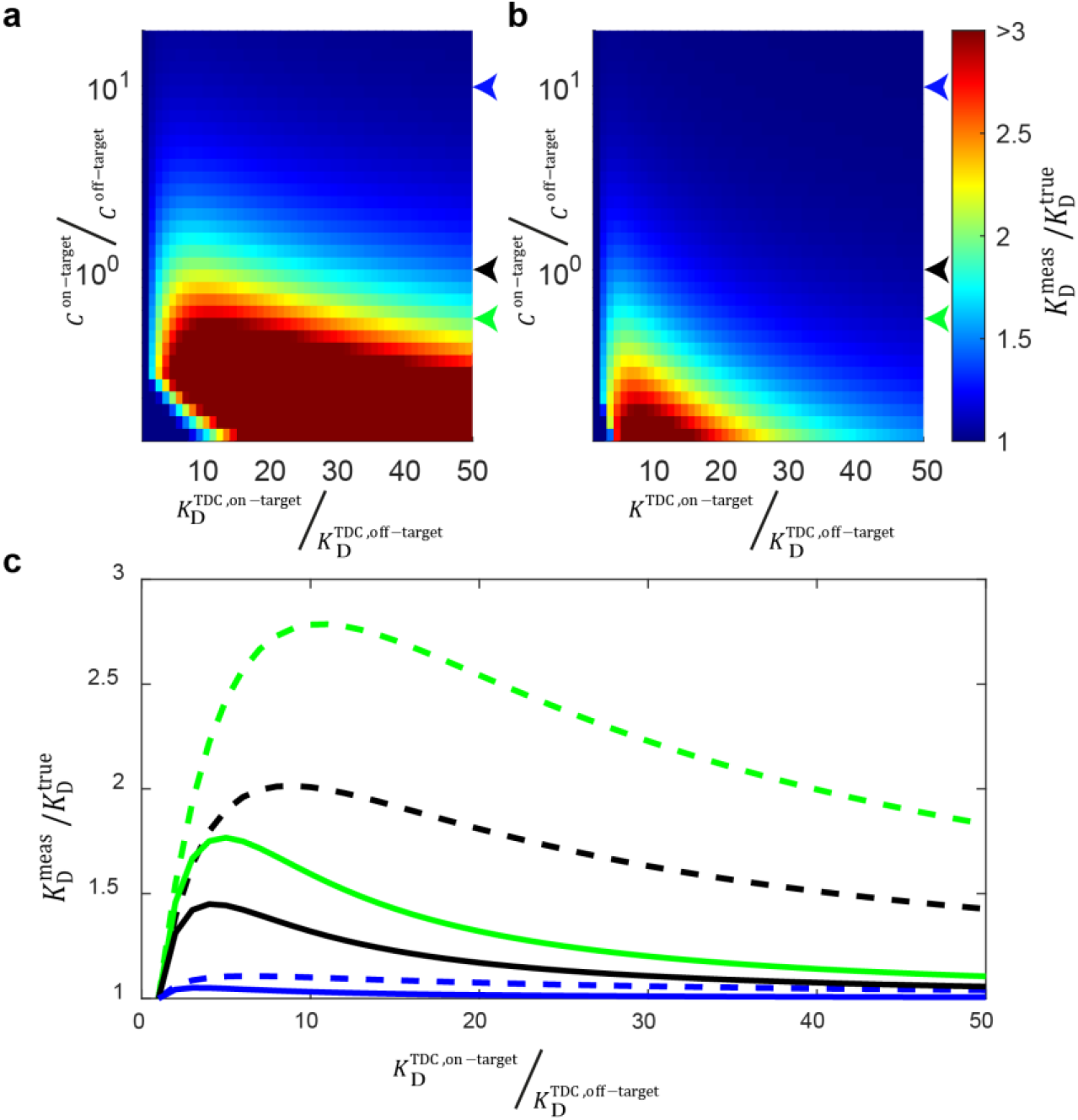
Simulated data for 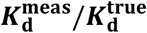 represented in surface plots. **(a&b)**The accuracy of the K_d,_ determination shown as 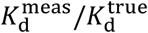 as a function of the concentration ratio *C*^on-target^/*C*^off-target^ between on- and off-target binding sites is displayed and off of the TDC specificity 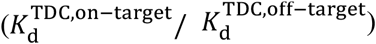 when **(a)** all binding events to the sensor surface are taken into account and **(b)** when the analysis is restricted to residence times longer than 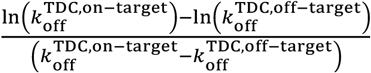 In **(a)** and **(b)**, the colour map indicates the ratio of the measured 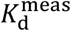 and the true 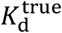. The arrows indicate the selected on- to off-target concentration ratios used in **(c)**, which shows cross sections extracted from **(a)** and **(b)** at on- to off-target concentration ratios of 0.5 (green), 1 (black), and 10 (blue), without (dashed line) and with (solid lines) kinetic selection of binding events.

An obvious requirement to the TDC is that it should be hydrophilic. Too lipophilic TDCs will non-specifically bind to the CDLs and cause rapid accumulation of a high number of CDLs at the sensor surface making distinguishing single liposomes impossible.

### Practical considerations for the novel dISA method

In addition to versatility, applicability, physiological accuracy, and flexibility of TDC selection, the time to establish a GPCR screening assay and the actual throughput of the measurements are important features of technologies utilized in drug development. Working protocols for methods based on detergent solubilization of GPCRs usually require substantial time to develop, while, in comparison, production of GPCR-containing cell membrane-derived vesicles is undoubtedly more straightforward and likely more broadly applicable to different types of membrane protein targets. Considering the throughput, a single dISA experiment takes less than 4 min. Since for each dISA experiment, a new microwell in a 384 sensor-well plate was used, no time for sensor-surface regeneration was needed. Combined with automated liquid handling and imaging, this enables achieving 384 kinetic measurements within 24 h.

Reagent consumption is another important parameter for compound-screening methods. It is particularly relevant when working with membrane protein targets that are demanding to express. Here, we used β_2_AR-CDL concentrations in the picomolar range for each inhibition experiment. Based on our experience, a single T-175 cell culture flask provides enough liposomes for >1000 single dISA experiments. Therefore, the dISA method is easily scalable and it is possible to produce enough target protein-containing CDLs for large screening campaigns, typically required for drug development, even with basic cell culture equipment.

In addition, the developed assay is generally applicable to any membrane protein, and thus serves as an attractive alternative to the many means that have been developed to overcome challenges related to the use of directed binding assays for membrane protein analysis, such as immobilization of detergent-solubilized membrane proteins^6^. The latter approach requires tedious assay development to adapt the membrane protein to a membrane-free environment^39^, and many membrane proteins change structure and lose function upon detergent solubilisation^14^. An alternative approach is to reconstitute detergent-solubilized membrane protein target into a natural lipid environment, such as liposomes^12^, or the so-called lipid nanodisks^9^. Even though these approaches allow to increase the surface coverage of GPCRs compared with e.g. immobilization of native cell membranes^9,64^, generation of surface coverage that yields sufficient signal-to-noise levels for kinetic profiling of low-molecular-weight compounds remains challenging^10^. All these limitations do not affect the here presented dISA method.

Finally, in any kinetic profiling study, it is the actual kinetics of the interaction between the target and the test compounds, that define the minimum time required for a single measurement. To overcome this bottleneck, the majority of label-free surface-based methods rely on different microfluidics-based protocols enabling running several measurements in parallel^1,2,65,66^, combined with regeneration protocols for multiple use of the same channel^67^. As mentioned above, the microplate format of the sensor plate allows to use a new microwell for each dISA experiment and abolishes thereby the need for sensor surface regenerations, which contributes to the short overall experiment time.

## Conclusions

We here presented a novel dISA methodology for full kinetic profiling of the interaction between suspended drug compounds and membrane-residing targets. We demonstrated the utility of dISA using β_2_AR, a major pharmaceutical target from the GPCR family. This was done by temporally following the transition between different interaction equilibria upon a change in the concentration of the inhibiting (screening) compound, *C*_inh_. We show that the rate of the transition between two such states is given by *k*_on_*C*_inh_ + *k*_off_, where the rate constants represent the interaction between the suspended drug compound and the membrane protein receptor contained in the liposomes. Further, since *K*_d_ = *k*_off_/*k*_on_ is also directly obtained from the degree of inhibition reached upon establishment of a new drug compound–induced quasi-equilibria^63^, full kinetic profiling can be obtained, as here demonstrated for the interaction between the agonist fenoterol and antagonist alprenolol to β_2_AR.

The assay operates in a total volume of less than 100 μL, with a concentration of the β_2_AR-containing liposomes in the low pM range, and has been designed for array-based readouts compatible with rapid screening. Because very low liposome concentrations are used, the assay is compatible with low- and high-affinity drugs, and offers a dynamic range that cannot be easily reached using conventional ISA because of sensitivity limitations^65^. In particular, this can be achieved by using a single TDC, which overcomes the need for different TDCs matched to the kinetic properties of the screened compounds, as is usually the case with conventional ISA^38^. Further, since the inhibition is governed by the interaction between the suspended drug compound and the membrane protein, the assay is applicable to drug compounds of any size, ranging from small fragments to large antibodies. In addition, unspecific binding of drug candidates to the liposome membrane does not greatly affect the measured kinetics. Of note, while we have demonstrated the dISA concept using cell-derived liposomes from CHO-K1 cells, numerous other methods exist for proteoliposome production, including cell-free expression, cell-derived vesicle formation^68^, and reconstitution into liposomes or nanodiscs^9^. These preparations are fully compatible with the dISA concept, making the method directly applicable beyond GPCRs screening. The array-based readout is also directly compatible with the growing use of DNA-encoded drug compound libraries^69^. Compounds from DNA-encoded libraries can be directly tethered to a sensor surface via their DNA label. The low concentration used and the high number of different sensor surfaces available will enable a rapid hit and target engagement validation for compounds identified in DNA-encoded library screens, followed by kinetic profiling of identified hits.

In summary, dISA offers a fast and reliable way to set up screening campaigns targeting GPCRs. The general applicability of this method to other target families and the fact that prior knowledge about the target is not needed should enable researchers to faster and more reliable target new GPCRs and deliver data of higher biological relevance for already well-known targets.

## Methods

### Cell line generation

A stable β_2_AR-overexpressing cell line was produced by transfecting CHO-K1 cells (ECACC) with a plasmid coding for the human β_2_AR with C-terminal GFP using lipofectamine 2000 (Invitrogen), according to the manufacturer’s instructions. To select stable clones, the cells were plated in a 60-mm dish and grown in “Ham’s F-12 Nutrient Mix, GlutaMAX™ Supplement” (Gibco), containing 10% v/v fetal bovine serum (FBS) (Gibco), and 600 μg/mL G418 reagent (Gibco) in a humidified atmosphere at 37 °C and 5% CO_2_. After single colonies became visible, cells from approximately 40 different colonies were aspirated with a 20 μL pipette and transferred to a 96-well clear bottom microplate. Transgene expression was assessed visually by fluorescence microscopy. Clones showing a high, homogenous, and stable expression were expanded and cryopreserved.

### CDV generation and purification

CDVs were generated as described elsewhere^41^, with some modifications. Briefly, CHO-K1 cells stably expressing the human β_2_AR were grown in a T175 flask until approximately 80–90% confluence in in Ham’s F-12 Nutrient Mix, GlutaMAX™ Supplement (Gibco), containing 10% v/v fetal bovine serum (FBS) (Gibco), and 600 μg/mL G418 reagent (Gibco). Before induction of CDV shedding, the cells were incubated for 2 h in Ham’s F-12 Nutrient Mix, GlutaMAX (Gibco) without serum. Prior to CDV shedding, 50 μL of Vybrant DiD reagent (Invitrogen) was added to the T175 cell culture flask, and the cells incubated for 20 min at room temperature in the dark. Subsequently, the cells were incubated for 30 min in, GlutaMAX (Gibco) medium supplemented with 10 μM/L cytochalasin B (ACROS Organics). The flask was shaken at 300 rpm at 37 °C during the incubation. The medium was collected in a 15 mL tube, and whole cells and debris were sedimented by centrifugation for 5 min at 700 × *g* and 4 °C. The supernatant was concentrated by passing through centrifugal filters with a 100 kDa molecular cut-off weight (MerckMillipore), to a volume of approximately 1 mL. The supernatant was then supplemented with protease inhibitor and extruded through a 100-nm pore filter using an Avanti Mini Extruder (Avanti Polar Lipids), seven times, to yield β_2_AR cell derived liposomes(β_2_AR-CDLs). The extruded supernatant was stored at 4 °C. The labelled β_2_AR-CDLs remained functional for several weeks when stored at 4 °C.

### Biosensor preparation

A cover glass (148 × 90 × 0.17 mm) was cleaned for 30 min in a solution of 1:1 30% ammonium hydroxide and 30% hydrogen peroxide at 90 °C. The cover glass was subsequently rinsed with water, dried, and bound to a bottomless 384-microtiter plate (Grainer Bioscience) with squared wells and 3.2mm base width. The surface was passivated by adding 25 μL of a 100:1 mixture of PLL-PEG/PLL(20)-g[3.5]-PEG(2)/PEG(3.4)- biotin(20%) (SuSos) solution to each well. The biosensor plate was incubated over night with the PLL-PEG/PLL-PEG-bio solution but incubation could be extended to several days. The plate was washed with HBS using a microplate washer, for a minimum dilution of the PLL-PEG/PLL-PEG-bio solution with HBS of 1:1,000,000. Then, a solution containing 1 μM of biotin-alprenolol (CellMosaic) and 1 μM Neutravidin (ThermoScientific) was prepared, and incubated for 30 min at room temperature. Subsequently, 25 μL of the solution was added to each well. The microplate was incubated at 16 °C and orbital shaking at 400 rpm for 2–4 h. After washing with HBS, and with a residual volume of 20 μL, the biosensor was ready to be used.

### Single-molecule platform

The main components of the single-molecule platform were a commercial TIRF microscope platform (Ti2e, Nikon) and an automated liquid-handling platform (modified OT2 robot, Opentrons). The microscope was equipped with a motorized XY-stage and a perfect focus system. For all measurements, the CFI Apochromat TIRF 60XC Oil, N.A. 1.49, was used. Images were acquired using a Hamamatsu ORCA-Fusion Digital CMOS camera (C14440-20UP). The microscope was controlled via NIS-Elements AR (Nikon). The OT2 robot was mechanically modified so it could be placed on top of the microscope and the pipettes could reach into the microplates mounted on the TI2-S-HW well plate holder (Nikon). The robot was equipped with two single-channel pipettes, P20 and P300. The robot was the leading system in the setup. The experimental protocols were written as Python scripts running on the OT2 integrated raspberry pi. The build commands were expanded by the necessary commands for the communication with the NIS-Elements AR software (Nikon).

### dISA experiment

Before the start of the experiment, the focus of the microscope was adjusted by adding few liposomes to a single well. After the focus was adjusted, the perfect focus system was activated. The fluorescence was excited using a 633 nM laser and the laser power was adjusted before an automated experiment run was started. The camera settings were 10 frames per s and 40 ms exposure time with standard read-out speed. Subsequently, the microscope was controlled by the liquid-handling platform. As the first step, 20 μL of the liposome-containing solution was added to a well and the data acquisition was initiated. After 20 s, 10 μL of a solution containing the ligand was added to the well and the solution was rapidly mixed. Solutions containing different concentrations of the ligand were prepared before the experiment in a 96-well plate and placed on the OT2 working deck. The OT2 robot delivered the solutions to the biosensor as per the defined protocol.

### Image analysis

To detect single liposomes, the local maxima were first detected by image dilation. All the detected local maxima were fitted using a 2D Gaussian function in GPUfit^70^. Particles were selected based on the amplitude and the sigma parameter of the 2D Gaussian fit. To differentiate newly arrived particles from newly detected particles and to supress unspecific binding events, particles were linked to trajectories, whereby the maximal allowed jump distance between frames was limited to 150 nm. Further, trajectories shorter than 1 s were discarded.

### Statistical analysis

#### Estimation of the confidence of fit parameters by non-parametric case resampling bootstrapping

To calculate the confidence interval of the inhibition level *β*_*eq*_and *k*_*obs*_, all binding events (*n* in total) that were observed in experiments where the same amount of test compound was added were pooled in a vector *x*_*i*_ = (*x*_1_ …, *x*_*n*_). Out of the vector *x*_*i*_ a new vector vector *x*_*i\**_ was generated by resampling n-times out of vector *x*_*i*_ with replacement. This was repeated 100 times. Each vector *x*_*i\**_ generated by resampling was analyzed as described in the theory section to calculate *β*_*eq*_and *k*_*obs*_. As error bars, the standard deviation of each fitted parameter was plotted.

## Acknowledgements

The authors thank Benjamin Krevet and Christian Pastucha for support with engineering and fitting the liquid handling platform to the microscope platform. We thank Stefan Geschwindner and Anders Gunnarsson for all their valuable discussions and input. We thank Catherine Kitts, Kees Oord and the whole Nikon team for brilliant support with hard- and software integration.

## Author contributions

TK and FH conceived the work. TK and VPZ contributed to the mathematical background of the approach. TK conducted the experiments and analysed the data. TK and FH discussed the data. TK and FH wrote the manuscript

## Data availability

