## Supplemental Information for "Single-molecule Dynamic In-Solution Inhibition Assay: A Method for Full Kinetic Profiling of Drug Candidate Binding to GPCRs in Native Membranes"

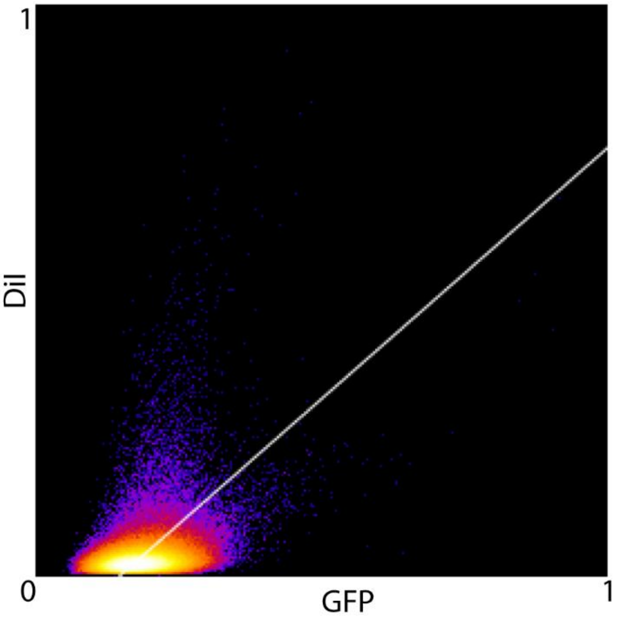


**Supplementary Figure 1 Colocalization analysis of DiI with $\beta_2$AR2-GFP.** 2D histogram of DiI vs GFP fluorescence from cell derived liposomes produced from DiI prelabeled cells. DiI and $\beta_2$AR2-GFP show little colocalization with a Pearson R value of 0.43 and a Li's ICQ value of 0.109.

### 1. Conditions for the applicability of the mean-field kinetic equations

Our analysis in the main text is based on the mean-field approximation (Eqs. 1-10) and implies uniform distribution of the concentration of the inhibiting compound and vesicles in solution. Under steady-state conditions, this is obviously the case. Under transient conditions, the assumption about uniform distribution should, however, be discussed and clarified from two perspectives.

First, any difference in the attachment and detachment rate of liposomes at the sensor surface will introduce a concentration gradient of the liposomes. The model we use do not take concentration gradients and diffusion limitation into account. Therefore, such gradients can, if not sufficiently small, in principle influence the observed transient kinetic regimes, and thus the kinetic analysis. The largest difference between the attachment and detachment rates of liposomes to the sensor surface is present directly after the solution with the $\beta_2$AR-CDLs is added to the microwell. The constant binding rate of the $\beta_2$AR-CDLs before addition of the inhibitor (at $t = t_{\mathrm{inh}}$) shows that no transient gradient exists in the solution Fig 4c, which suggests that the diffusional motion of the liposomes counteracts any transient gradient, and that the transient gradients are sufficiently small for the concentration of liposomes close to the surface to be quasi-equal to average total concentration of liposomes $C_{\mathrm{tot}}$.

Furthermore, the ability to detect, count and track single liposomes makes it possible to detect relative changes in the concentration of the $\beta_2$AR-CDLs. It is in this context also worth noting that the number of liposomes that can be tracked is a subfraction of the total liposomes interacting with the surface, since only liposomes that stay long enough in the focal volume can be detected and tracked. Still, this number is proportional to the total liposome concentration close to the surface and can therefore is used as relative measure of concentration changes. The number of free vesicles shows no significant variation during an inhibition experiment, except for stochastic fluctuation and the dilution induced by the inhibitor injection (Supplementary Figure 2).

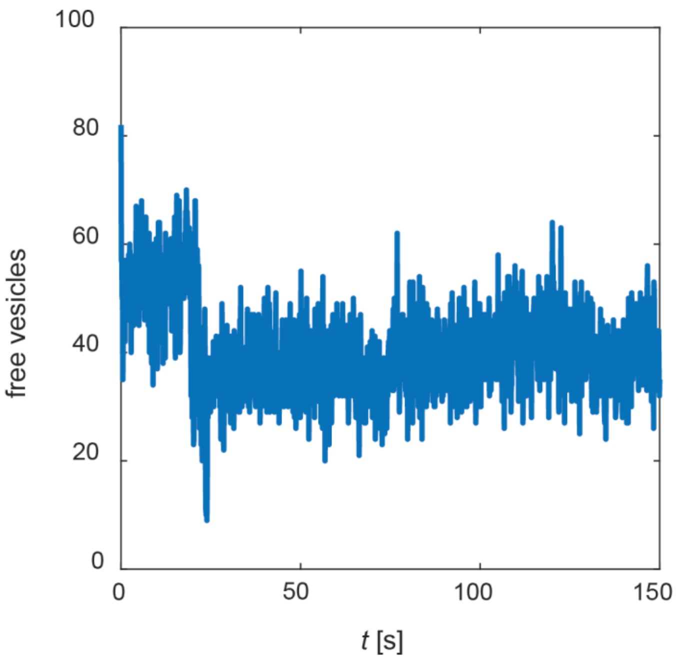


**Supplementary Figure 2 Absence of local concentration fluctuations at the sensor surface.** Number of free vesicles detected during a representative inhibition experiment.

Second, the mixing by repetitive pipetting with the inhibitor solution could induce a fluid flow in the microwell. The ability to track single liposomes allows to time resolve any liquid flow in the solution, which makes possible to track changes in the average velocity of the liposomes over time. Supplementary Figure 3 shows the average velocity of the liposomes over time, including two repetitive pipetting steps (at $t$ = 20 s), demonstrating that the average velocity of the liposomes returns to average diffusional velocity within 0.8 s, which is significantly shorter than the transition time from which the kinetics is extracted.

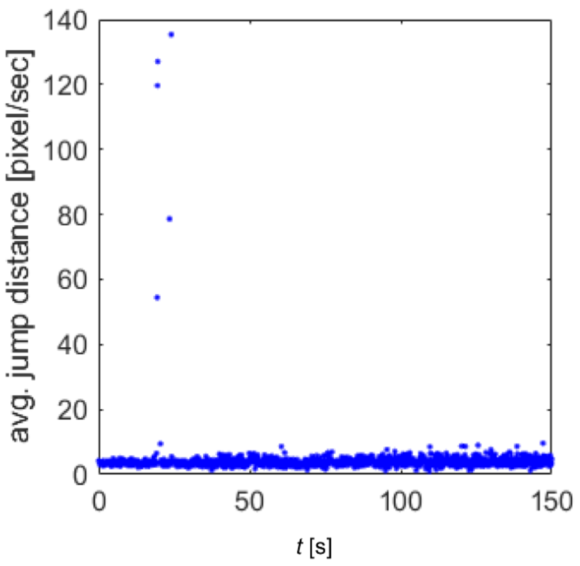


**Supplementary Figure 3 Rapid decay of forced fluid flow during mixing.** The mean velocity of the liposomes after mixing of the $\beta_2$AR-CDL already in the test well with a buffer solution containing the inhibitor at ~ $t$ = 20 s. The velocity increases during 8 frames, corresponding to 0.8 s.

This finding is in agreement with the following theoretical considerations. After mixing ($t > t_{\text{inh}}$), the liquid distribution in the measurement cell is described by the Navier-Stokes equation for velocity distribution in an incompressible liquid (see Eq. (24.2) in Ref. [1]

$$\frac{\partial \mathbf{v}}{\partial t} = -\frac{1}{\rho}\mathbf{grad}p + \nu\Delta\mathbf{v} \tag{S1}$$

where $p$ is pressure, $\rho$ is the density, and $\nu$ = 0.010 cm$^2$/s is the solution (water) kinematic viscosity. According to the equation, the maximum timescale of relaxation of this movement in the solution is given by

$$\tau_{\text{rel}} = \frac{L^2}{\nu} \tag{S2}$$

where $L$ is the length scale characterizing the measurement cell with solution (roughly, one half of the minimum cavity size). For applicability of Eqs. 1-10, this timescale should obviously be shorter than the timescale, $\tau_{\text{rel}}$, characterizing the transient period of the kinetics. In our case, we have $L \cong$ 0.15 cm, and accordingly $\tau_{\text{rel}}$ = 2.25s, which is significantly shorter than the transition time.

### 2. Selection of binding events by residence time at the sensor surface

The ability to select binding events based on their residence time at the sensor surface is a feature of this method that can be used to discriminate on- and off-target binding events. Typically, the difference between on- and off-target binding is described by the difference of the respective dissociation's constants $K_d$. For the discrimination of on- and off-target binding, the binding kinetics as described in the following are decisive. The residence time at the sensor surface is determined by the respective off-rates of the on- and off-target binding. The number of binding events depends on the respective association rates of the on- and off-target binding. For this analysis, we assume that the TDC binds next to its target to a single type of off-target binding site, with the dissociation constant for the on-target binding being given by

$$K_{\mathrm{d}}^{\mathrm{TDC,on-target}} = \frac{k_{\mathrm{off}}^{\mathrm{TDC,on-target}}}{k_{\mathrm{on}}^{\mathrm{TDC,on-target}}} \tag{S4}$$

while for the off-target binding, it is given by:

$$K_{\mathrm{d}}^{\mathrm{TDC,off-target}} = \frac{k_{\mathrm{off}}^{\mathrm{TDC,off-target}}}{k_{\mathrm{on}}^{\mathrm{TDC,off-target}}} \tag{S5}$$

Furthermore, we assume that the dissociation rate and association rate of the off-target binding contribute to the same extend to the dissociation constant of the off-target binding. Therefore:

$$k_{\mathrm{off}}^{\mathrm{TDC,on-target}} = \sqrt{\varphi} k_{\mathrm{off}}^{\mathrm{TDC,off-target}} \tag{S6}$$

where:

$$\varphi = \frac{K_{\mathrm{d}}^{\mathrm{TDC,on-target}}}{K_{\mathrm{d}}^{\mathrm{TDC,off-target}}} \tag{S7}$$

This means that $k_{\mathrm{off}}^{\mathrm{TDC,off-target}}$ can be expressed as:

$$k_{\mathrm{off}}^{\mathrm{TDC,off-target}} = \frac{k_{\mathrm{off}}^{\mathrm{TDC,on-target}}}{\sqrt{\varphi}} \tag{S8}$$

The residence time threshold for selection of binding events can be arbitrarily set, but there is a trade-off between suppressing the off-target binding and abolishing falsely on-target binding events as can be seen in Supplementary Figure 4a. As threshold we use here the timepoint where the absolute difference between the probability density distribution of the residence time of the on- and off-target binding has its maximum (Supplementary Figure 4b). The difference between the probability density distribution of the residence time of the on- and off-target is

$$\Delta(t) = e^{-k_{\mathrm{off}}^{\mathrm{TDC,on-target}} t} - e^{-k_{\mathrm{off}}^{\mathrm{TDC,off-target}} t} \tag{S9}$$

This function has its maximum at

$$t_{lim} = \frac{\ln\left(k_{\mathrm{off}}^{\mathrm{TDC,on-target}}\right) - \ln\left(k_{\mathrm{off}}^{\mathrm{TDC,off-target}}\right)}{\left(k_{\mathrm{off}}^{\mathrm{TDC,on-target}} - k_{\mathrm{off}}^{\mathrm{TDC,off-target}}\right)} \tag{S10}$$

This means that the possibility of discriminating between on- and off-target binding is only dependent on the ratio $\varphi$ (Eq. S4) of the on- and off-target dissociation constants.

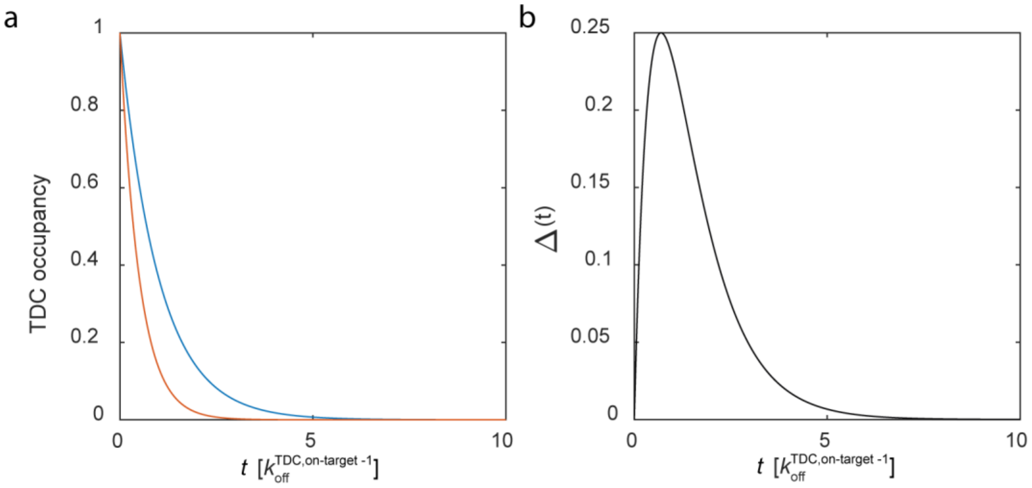


**Supplementary Figure 4 Differentiation of on- and off target TDC binding events by residence time. (a)** Exponential residence time distribution of the TDC on-target binding (blue) and two times faster dissociation of TDC off-target binding(red) **(b)** Difference $\Delta(t)$ of on- and off-target occupancy of the TDC.

Defining the discrimination for on- and off-target binding by $t_{lim}$ into account the convoluted sensorgram of on- and off-target binging were calculated and fitted as in the theory section of the main text described by Eq. 9. The fitted dissociation constant was $K_\mathrm{d}^\mathrm{meas}$ and the maximum theoretical reachable accuracy of measured $K_\mathrm{d}^\mathrm{meas}$ is described by the ratio $K_\mathrm{d}^\mathrm{meas}/K_\mathrm{d}^\mathrm{true}$ (see Fig 6, main text).

1. Landau, L. D. & Lifshitz, E. M. *Fluid Mechanics*. (Elsevier, 1987). doi:10.1016/C2013-0-03799-1.
